# Solving the Variable-Particle Size Quandary of Bone Mills. Development of an Automated Milling System to Generate Graft Particles of Definite Sizes

**DOI:** 10.1101/2023.01.13.523924

**Authors:** R Viswa Chandra

**Affiliations:** SVS INSTITUTE OF DENTAL SCIENCES

**Keywords:** Bone Grafting, Bone Mill, Hydroxyapatite

## Abstract

There are situations where block grafts have to be milled to convert them into particulate grafts of definite sizes. The objectives of this study were to evaluate the quality of graft particles generated in two sizes from a custom-built automated milling system (AMS) and their biocompatibility in an animal model. A Monetite block was milled in an AMS to generate small (SS group; 0.5-0.8 mm) and medium size (MS group; 1.0-1.2 mm) particles. Measures of particle count, Feret’s diameter (dF), particle distribution and size were recorded. Biocompatibility of particles was tested in a rabbit tibial defect model. The average particle size was significantly smaller in the SS group than the MS group (0.68±0.39 *vs* 1.10±0. 79 mm; *p*≤*0*.*001*). There were significant to highly significant differences between SS and MS groups in measures of particle count (*p*≤*0*.*001)*, dF *(p=0*.*02)* and size (*p*≤*0*.*001)*. SS and MS groups had maximum percentage of particles in the 0.6-1mm (71%) and >1mm (70%) ranges respectively. The mineralized tissue volumes across SS, MS when compared to an autogenous block were 68.92±35.66%, 66.29±29.21% and 89.83±19.91% (*p=0*.*003)* respectively. The device was able to generate small and medium-size graft particles which were distinct from each other.

## INTRODUCTION

Harvesting and milling of bone substitutes is an instrument-dependent procedure.^1-4^ Different techniques yield grafts of varying particle sizes; at around 3-4 mm, particles harvested by means of hand chisels are the largest and most inconsistent whereas graft particles obtained by use of high and low-speed burs are within the range 0.3-0.5 mm.^5-7^ The bone-forming ability is again dependent on the particle size. Depending on the size, location and nature of the defect, particle sizes ranging from 150-600 µm to 1-2mm are considered essential for vascularization, graft-incorporation and subsequent bone formation.^5-8^

The particle size is however locked to the instrument used to obtain it.^1,3,5,7,9^ Rotary instruments can harvest graft particles of varying sizes; from 250 to 500 µm,^3,4^ 500–1000 µm^4,7,8^ or beyond 1000 µm^4,7^. There are situations prevailing in which, novel materials such as tooth-derived bone grafts,^10^ or grafts such as autogenous bone blocks,^5,7^ block allografts,^11^ xenogeneic or alloplastic blocks^6^ have to be subsequently milled to convert them into particulate grafts. The optimum particle size is 75-500 µm, 0.9 to 1.7 mm, 0.75-1 mm and 250 to 650 μm for tooth-derived grafts,^10^ block allografts,^11^ xenogeneic grafts and alloplastic blocks^6^ respectively. To generate particulate grafts in such range of sizes can be accomplished only with a wide range of armamentarium which may not be feasible practically or economically and a device to generate grafts of different particle sizes will be a welcome addition to a dentist’s arsenal. A bone mill should ideally have the following features;^12^ 1. It should preferably be power driven with an ability to generate particles of variable sizes on different materials including dentine, bone and alloplastic materials.^12,13^ 2. The graft particles should be generated in a shorter time at lower rotation speeds. (minimum-pass cutting) to prevent thermal damage to the material being processed.^3,7,14^ and 3. The device should be of simple construction to facilitate easy cleaning and decontamination.

In this context, the specifications mentioned above were taken into consideration in the design and development of an automated milling system (AMS) capable of generating particles of two distinct sizes. This study aimed to investigate the morphometric qualities of particles obtained and to assess the biocompatibility of these graft particles in an animal model.

## MATERIALS AND METHODS

An automated milling system *(Figure 1a)* was constructed in surgical-grade stainless steel (conforming to IS 5583:1970 and IS 5589:1970, Bureau of Indian Standards) to generate graft particles between 0.5-0.8 mm and 1.0-1.2 mm. Briefly, the device has two rings (diameter: 50 mm; distance: 3 mm) of flat burrs^12,14^ mounted perpendicularly to the long axis and varying the RPM enables to generate small (SS group; 0.5-0.8 mm) and medium size (MS group; 1.0-1.2 mm) *(Figure 1b)* particles. The primary objectives of this study were 1) To evaluate the quality of graft particles generated in two sizes 2) Because grinding^9,12,14^ may affect the quality and compatibility of graft particles with human tissues, their biocompatibility was evaluated in an animal model.^9^ Both the objectives had specific methodologies that were conducted concurrently. Each methodology had two groups (SS & MS) based on the size of the particles obtained.

**Figure 1a:**
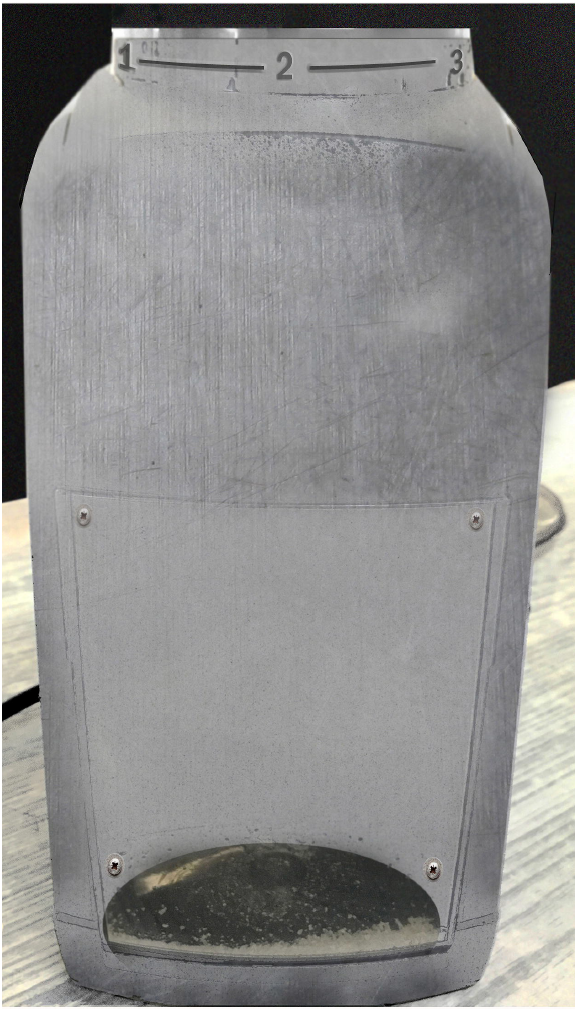
An automated milling system (AMS) to generate graft particles between 0.5-0.8 mm and 1.0-1.2 mm.

**Figure 1b:**
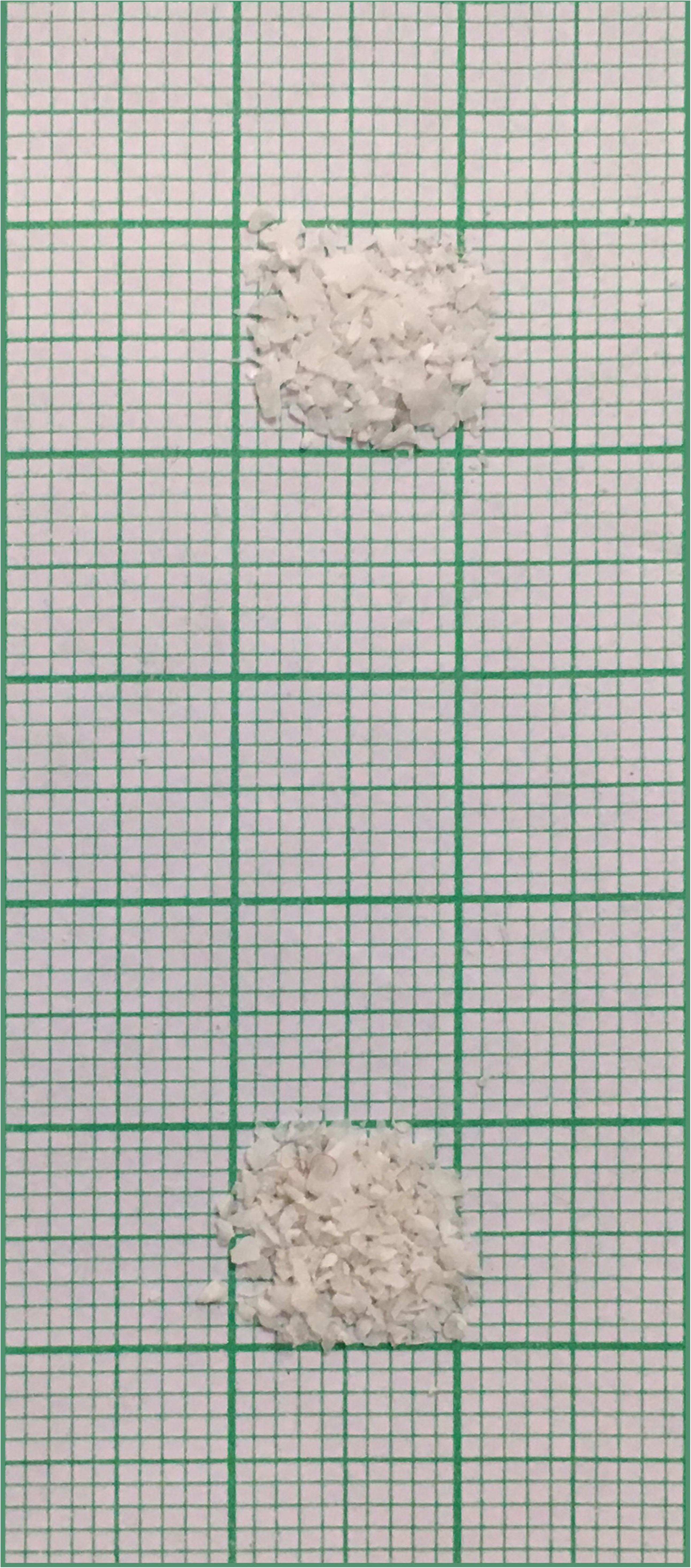
Graft particles between 0.5-0.8 mm *(bottom)* and 1.0-1.2 mm *(top)*.

**Figure 1c:**
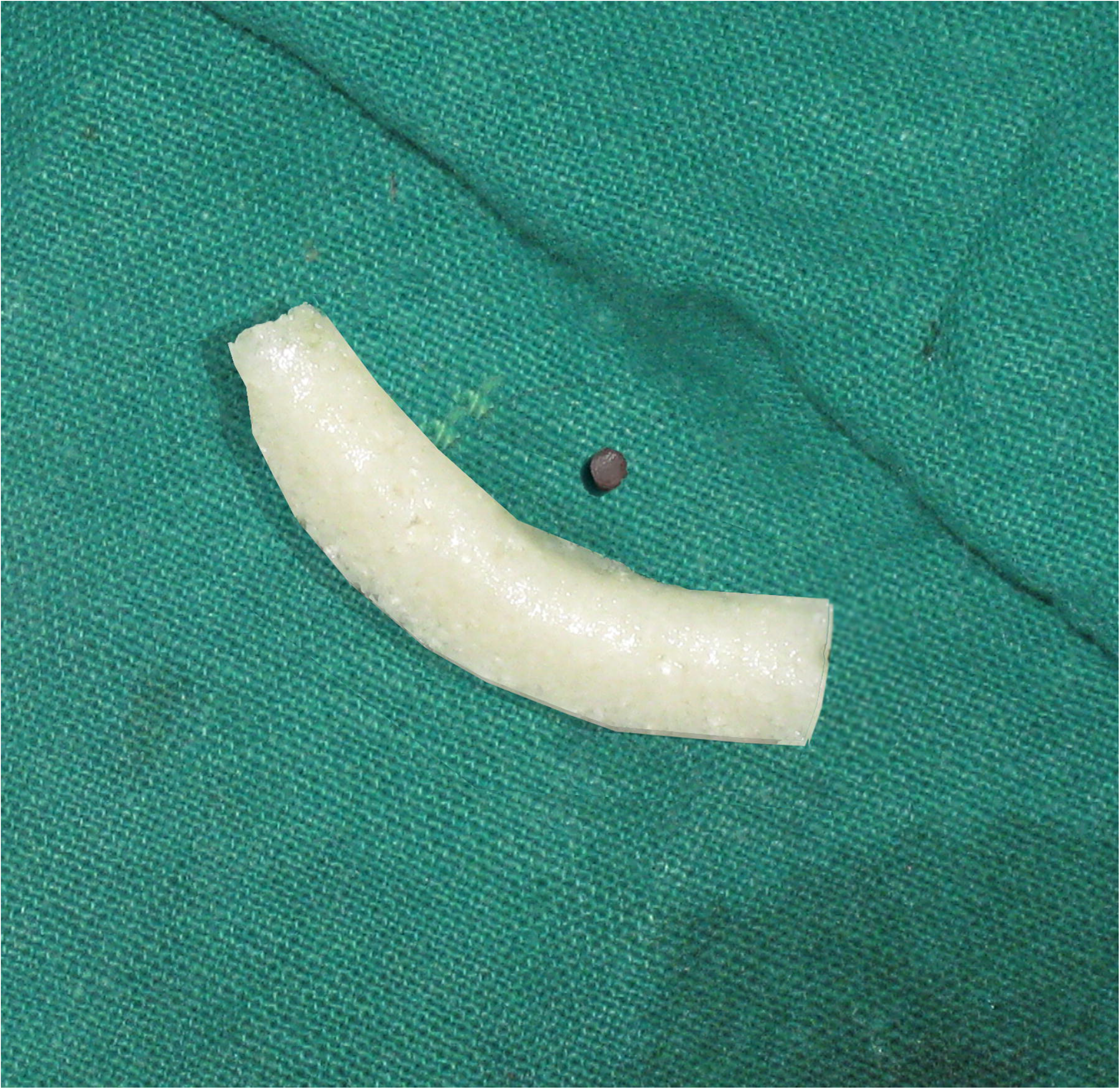
The automated milling system was used to cut the Monetite block into sizes corresponding to SS and MS groups respectively. With it is an autogenous core extracted from a different location which acted as a positive control.

### Morphometrical analysis of graft particles

#### Sample Size and Material Preparation

A minimum sample size of 40 per group was needed to detect a particle size difference of 0.1 mm when the power of the test is 0.80 at a significance level of 0.05. Accordingly, a material stronger than bone in the form of curved Zn-substituted Monetite block^15,16^ (density: 2.3-2.9 g/cm;^16^ moisture: 2.12%; *Figure 1c*) was used as a test material to allow for comparative tests after processing it in the AMS. The material was moistened and then placed *in toto* into the device to generate particles at settings 1 (500 RPM for 30s) or 2 (900 RPM for 30s) to generate MS and SS respectively. The obtained material was spread on a medical grade graph paper (MicroMed, Chennai, India) to create 50 “squares” of 1*1 sq. cm^2^ per group *(Figure 1b)*. Further analysis was done sequentially as follows.

#### Particle Count and Feret’s diameter (dF) estimation

Standardised photographs of all the squares with graft particles were taken with a macro lens from 100 mm at 1:1 magnification. Particle count *(Figure 2)*, size distribution and the Feret’s diameter (dF), which is the longest distance between any two points for a particle were measured with an image processing program (ImageJ®, LOCI at the University of Wisconsin-Madison, USA). Briefly, it was done as follows;^17,18^ After scaling (to convert pixels into millimetres) and thresholding, in the ‘set measurements box’, Feret’s diameter was specified and the number of particles from 0-∞ and with circularity of 0.4 were counted. The distribution of particles based on dF value was also calculated from the output *(Figure 3)*.

**Figure 2.**
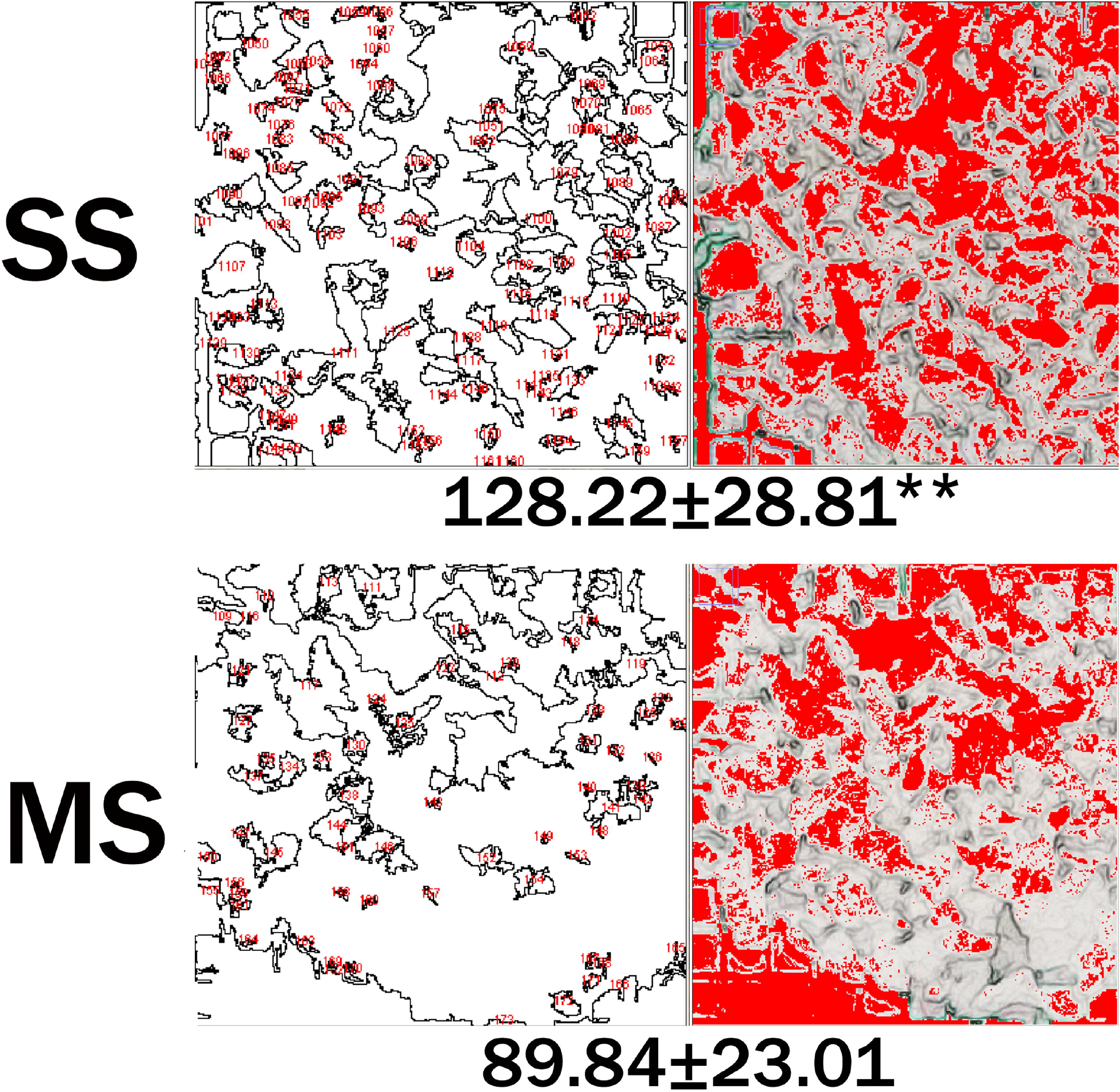
The particle count/cm^2^ showed a highly significant *(p*≤*0*.*001**)* difference between the SS and MS groups.

**Figure 3.**
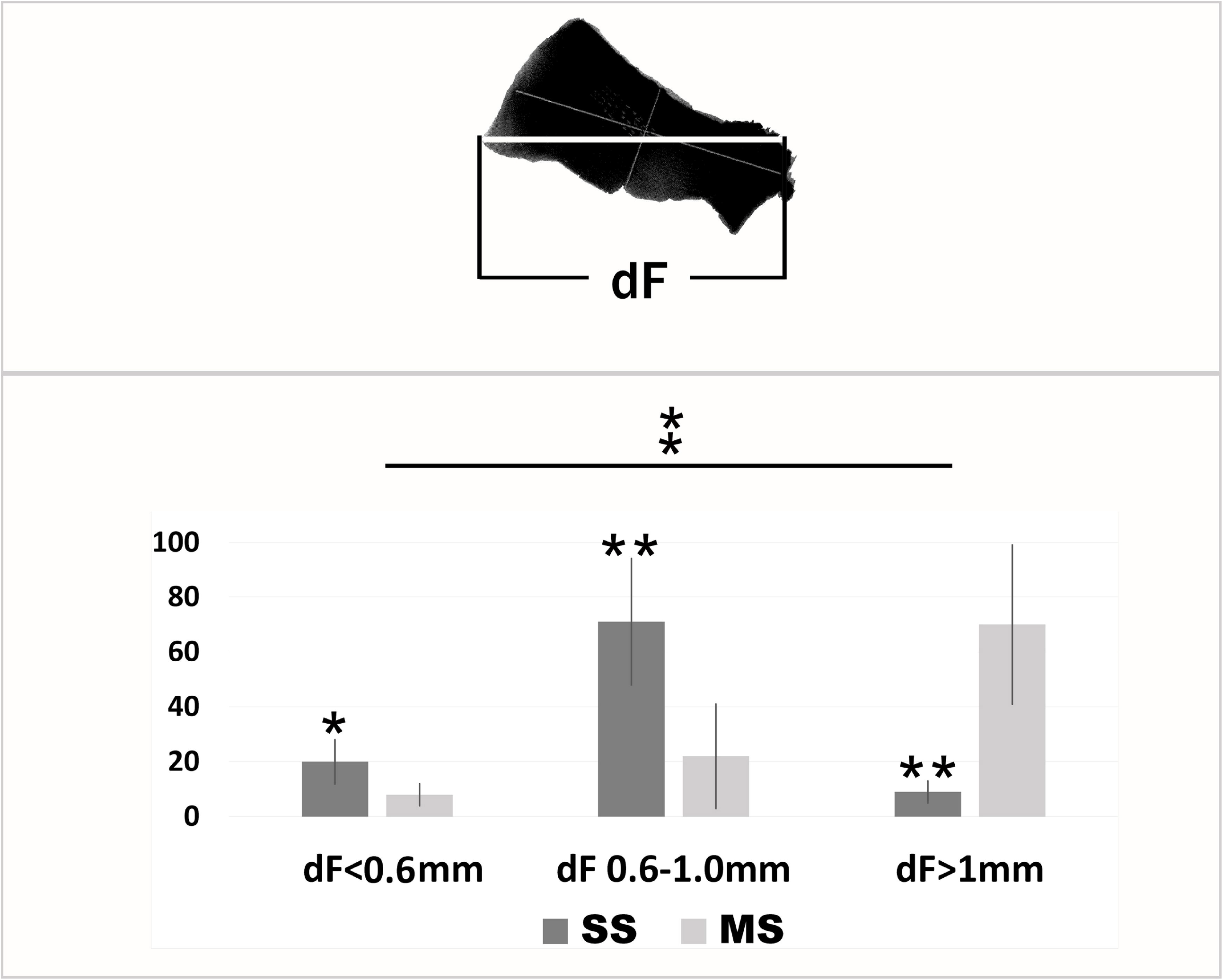
Feret’s diameter (dF) is the longest distance between any two points for a particle. The distribution of the particles based on dF seem to match with the size of the particles as there seems to be no overlap in the particle range. There were significant to highly significant intragroup (SS: *p*≤*0*.*001*^□^ and MS: *p*≤*0*.*001*^□^) and intergroup differences (<0.6: *p=0*.*01\**; 0.6-1.0mm: *p*≤*0*.*001\*\**; >1mm: *p*≤*0*.*001\*\**) at each range.

#### Particle Size estimation

Using a 6 mm micro spatula, 3 random scoops of particles per square in every group were transferred onto a Petri dish for particle size estimation using a Stereomicroscope (VISION^®^, Vision Micro Systems, Kolkata, India).^18^ Photographs of 20 particles per Petri dish at 1280 × 980 pixels (10X magnification) were taken. A calibration of 2µ/pixel was set and the lengths and the widths of the particles were measured. The average of length and depth was defined as the average particle size (APS). The length and breadth of the largest particle from one square was defined as the maximum length (ML) and maximum width (MW) *(Figure 4)*.

**Figure 4.**
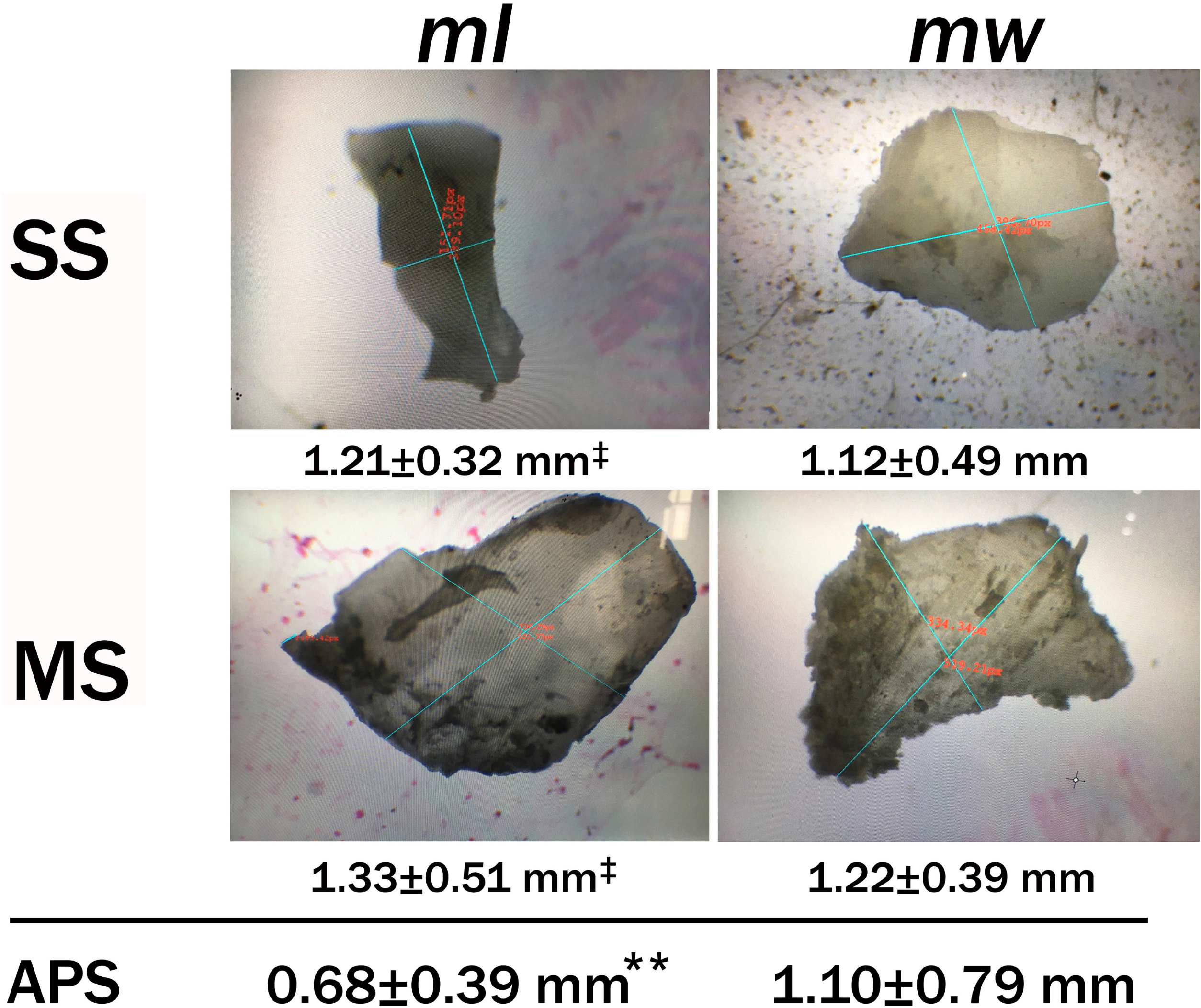
Though the particles were not spherical, there was no significant difference between mean length (ML) and mean width (MW) in both the groups (SS: *p=0*.*09*^‡^ and MS: *p=0*.*06*^‡^). The average particle size (APS) was significantly smaller in the SS group than the MS group (*p*≤*0*.*001\*\**).

### Histomorphometric analysis of the Graft Particles

#### Sample size, participants and interventions

To have an 80% chance (*β* error) of detecting a significant (two-sided 5% level) and a largest difference of 30 between groups with a standard deviation of 1, 20 sites were required. Accordingly, 10 healthy New Zealand rabbits (age: 4.20±1.1 months; mean weight: 3.45±0.65 kgs) were included in the study. Animals were selected after approval from the Institutional Animal Ethics Committee *(SVSDC/2017-6)* and the study was done in an accredited animal lab. Sedation and anaesthesia were induced through an intramuscular (IM) injection of Xylometazoline (Nazotic P© Biotic Healthcare, Panchkula, India; 3-5mg/kg) and Ketamine (Hypnoket©, CBC Pharma, Mumbai, India; 35-40mg/kg). The anteromedial aspect of the tibia was exposed by a 5cm-long longitudinal anterior incision. A 4mm trephine (Komet©, Komet Dental, Ahmedabad, India) was used to create three osseous cavities at a distance of 4 mm to each other and the autogenous cores were retained *(Figure 5a)*. From the proximal to distal tibia on both sides, each cavity was filled with medium (MS) & small size (SS) particles along with an autogenous core extracted from a different location which acted as a positive control (C) *(Figure 1c, 5b)*. Thus, each animal yielded two sites for each group (SM, MM & C; 20 sites/group, 60 sites in total). The soft tissue was repositioned and layer wise closure was done.

**Figure 5a:**
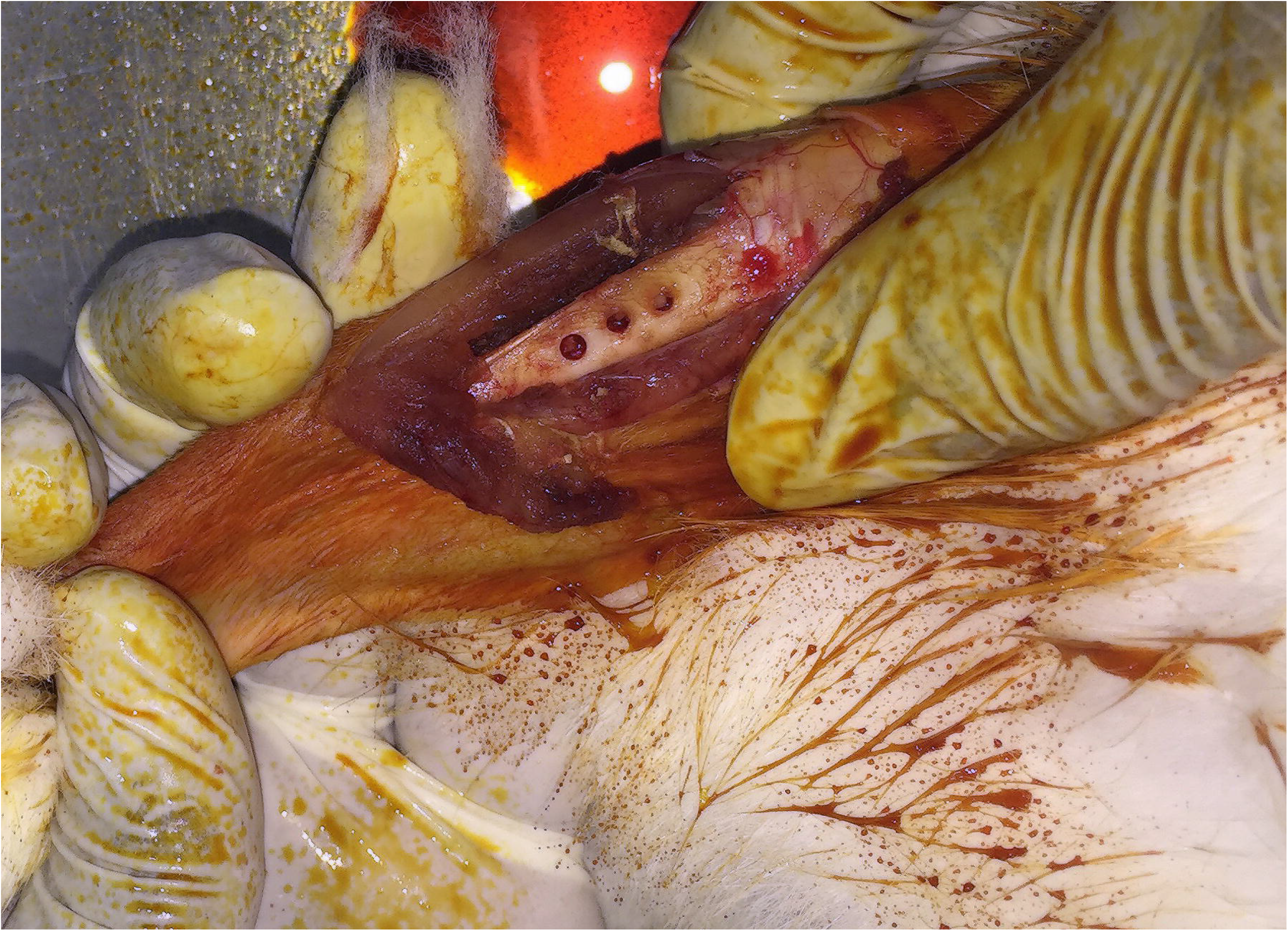
The anteromedial aspect of the tibia was exposed by a 5 cm long longitudinal anterior incision.

**Figure 5b:**
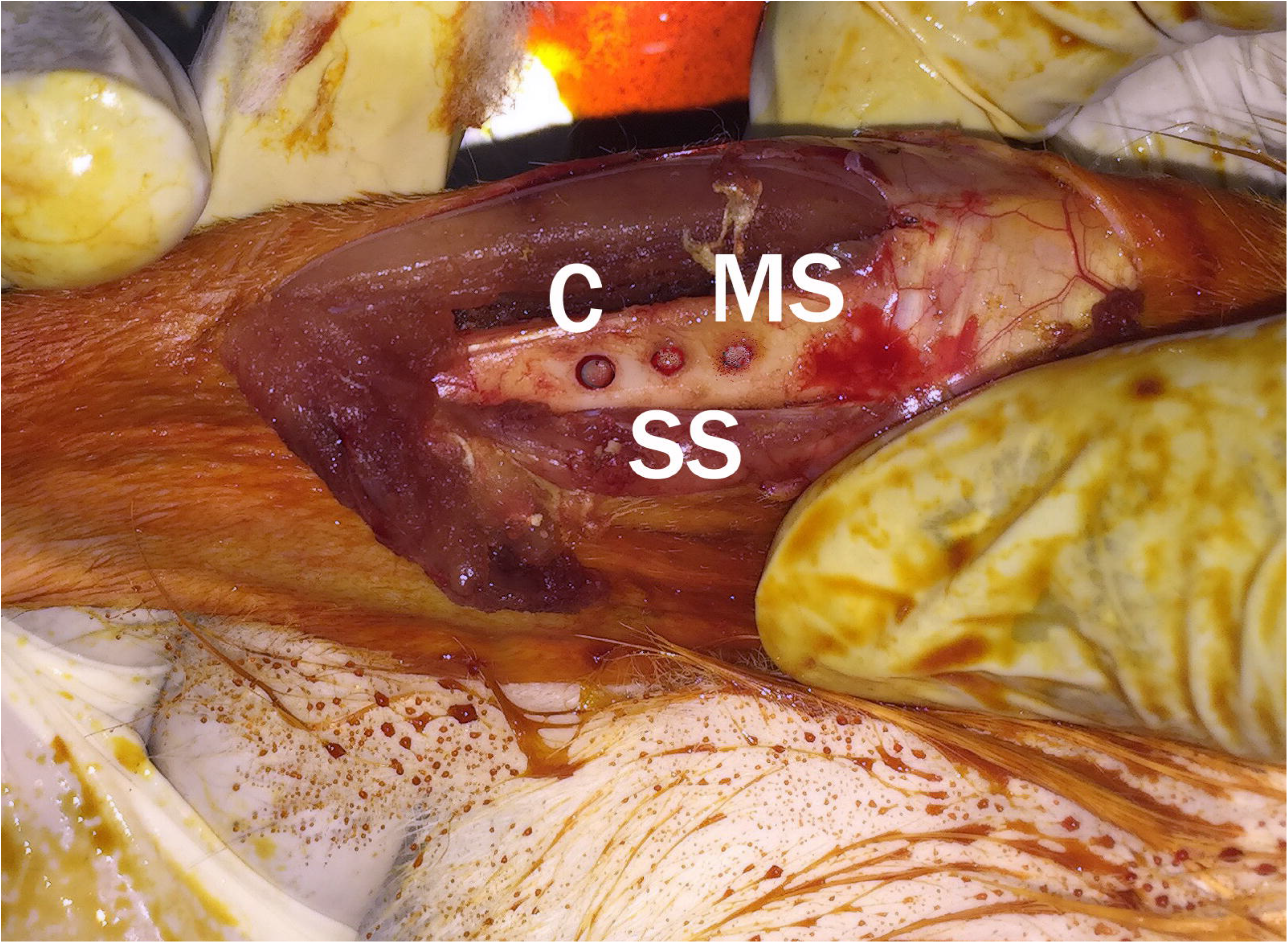
From the proximal to distal tibia, each cavity was filled with medium (MS) and small size (SS) particles along with an autogenous core extracted from a different location which acted as a positive control (C).

#### Tissue processing and histological analysis

2 months after the procedure, all rabbits were sacrificed with an appropriate dose of intravenous Ketamine and the tibiae were extracted. After dehydration and demineralization, five semi-serial longitudinal sections (5µ thick) from each of the twenty tibiae were subjected to routine haematoxylin-eosin staining for light microscopy analysis. From operative sites of each section, ten regions of interest (ROIs; *Figure 6*) per slide were imaged (Olympus BX53® microscope, DSS Group, New Delhi, India) at 40X magnification. The mineralized tissue volume (MTV) was assessed as follows;^15^ The images were opened in ImageJ®, binarized^17,18^ and areas of mineralized tissue/ bone were selected. Selecting Analyze>Measure quantifies the mineralized tissue volume (MTV) of the slide which is derived by the formula (mineralized tissue area/total area) *100. The MTV of a group was defined as the average of MTV values for 10 ROIs per slide for each section within a group.

**Figure 6.**
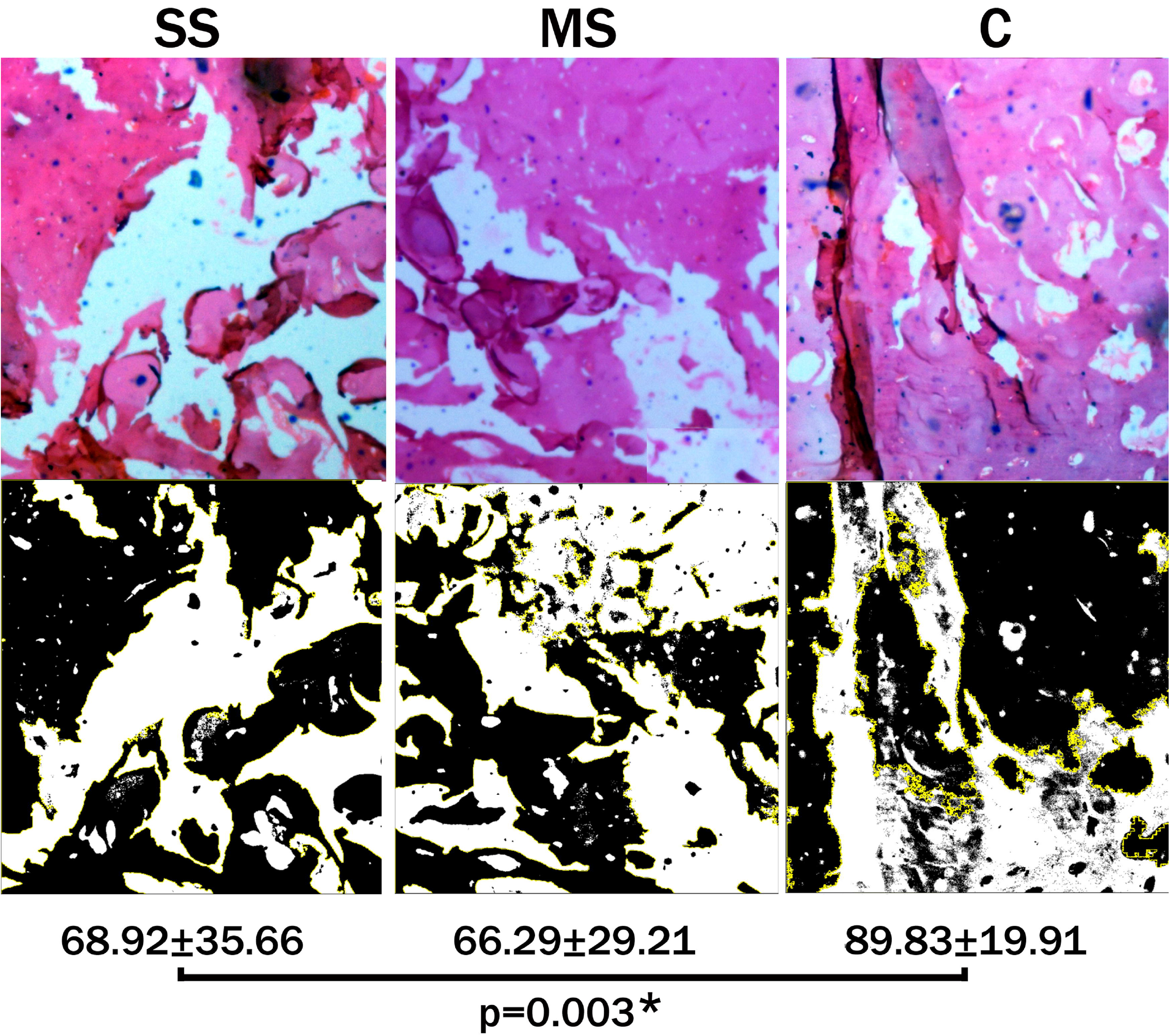
The images were opened in ImageJ® *(top row)*, binarized *(bottom row)* and the mineralized tissue volume (MTV) was measured. There was a significant intergroup difference among MTV values across SS, MS and C groups at 3-months *(p=0*.*003*)*.

### Statistical Analysis

Data was analyzed by using Prism8^®^ (GraphPad Software, La Jolla, USA). Intragroup comparison was performed by using ANOVA followed by multiple comparisons using Bonferroni correction. One-way ANOVA followed by the post hoc test was used for intragroup and intergroup comparisons. A p≤0.001 was considered as highly significant, p≤0.05 as significant and p>0.05 as non-significant.

## RESULTS

Across both groups (SS & MS), 100 ‘squares’ of particulate matter were evaluated for particle count, Feret’s diameter (dF) and particle distribution estimations. 2000 particles were imaged for particle size and average particle size (APS) estimation; furthermore, 100 particles were specifically imaged to derive maximum length (ML) and maximum width (MW). Histomorphometric analysis was performed in rabbits; data from one animal, which sustained a pathologic fracture within a week after surgery, was not included in the final analysis. Hence, mineralized tissue volume (MTV) was calculated for 18 sites in each group (SS, MS & C).

### Morphometrical analysis of graft particles

The particle count/cm^2^ showed a highly significant *(p*≤*0*.*001)* difference between the SS (128.22±28.81) and MS (89.84±23.01) groups *(Figure 2)*. There was a significant difference (*p=0*.*02*) in dF values between both groups as well (1.29±0.39 & 1.38±0.67 mm in SS and MS groups respectively). SS and MS groups had maximum percentage of particles in the 0.6-1mm (71%) and >1mm (70%) ranges respectively *(Figure 3)*. The distribution of the particles based on dF seems to match with the size of the particles as there seems to be no overlap in the particle range as evidenced by significant to highly significant intragroup (SS: *p*≤*0*.*001*and MS: *p*≤*0*.*001*) and intergroup differences (<0.6: *p=0*.*01*; 0.6-1.0mm: *p*≤*0*.*001*; >1mm: *p*≤*0*.*001*) at each range. Though the particles were not spherical, there was no significant difference between ML and MW in both the groups (SS: *p=0*.*09* and MS: *p=0*.*06*). The APS was significantly smaller in the SS group than the MS group (0.68±0.39 *vs* 1.10±0. 79 mm; *p*≤*0*.*001*) *(Figure 4)*.

#### Histological analysis

The MTV values across SS, MS and C groups were 68.92±35.66, 66.29±29.21 &89.83±19.91 respectively. There was a significant intergroup difference among these groups at 2-months *(p=0*.*003) (Figure 6)*.

## DISCUSSION

Hand-powered mills with crushing^19^ or cutting surfaces^20^ are popular and have been studied extensively. Manual bone mills produce larger-sized^20,21^ and irregularly shaped graft particles;^22^ their ability to form high-quality bone shows contradictory or conflicting results in various studies.^4,7,8,20-22^ In contrast, powered bone mills can produce particles of various sizes^4,8,10,12,13^ and can consistently produce smaller sized graft particles within a range considered as optimum for osteogenesis.^4,8,21,23^ However, limited availability, cost and narrow indications for usage have hampered their development. An automated milling system can be designed to generate graft particles of various sizes from block allografts, xenogeneic substitutes and alloplastic blocks as well.^10,11^

The prototype device was able to generate small (SS) and medium-size (MS) graft particles which were distinct from each other in measures of particle count, Feret’s diameter (dF), particle distribution and size. From cortical bone, Fucini et al^4^ were able to generate particle sizes of 250-500 μm and 850-1,000 μm using an analytic mill and Cornu et al^23^ were able to obtain particle sizes ranging from 0.2 to 2.9 mm from femoral heads by using a Retsch mill. Using an unspecified method of grinding, Nam et al^24^ were able to produce demineralized dentin matrix in two sizes, 0.25∼1.0 mm and 1.0∼2.0 mm. Interestingly, the analytic and Retsch mills use blades like ‘double beater’ or ‘cutting bars’ to process grafts at higher speeds (1500-28000 RPM). The device in this study has two rings of flat burrs; the uncut Monetite block goes through whole width of two burrs resulting in efficient cutting at lower speeds. ^12,14^

The particle count/cm^2^ and dF values showed significant differences between the SS and MS groups. It can be stated that both groups had different densities as in this context, particle count is a measurement of the number of particles in a spatial unit (1 cm^2^).^17,18^ The ability of a device to produce graft particles at varying densities and sizes is an important attribute. Smaller particles at higher densities show greater surface area which theoretically increase the available surface area for osteogenesis.^4,5,8,25,26^ Conversely, larger particles at lower densities are more resistant to resorption and micromovement^8,23^ and may show bone formation comparable to smaller-sized grafts.^4,21^ Particle distribution analysis performed after milling showed that the SS and MS groups had maximum percentage of particles in the 0.6-1mm (71%) and >1mm (70%) ranges respectively. The average particle size was significantly smaller in the SS group than the MS group and the distribution stayed true to the original particle sizes with no significant shift onto other size ranges. A trephine can be considered as a single-ended miniaturized version of a flat-burr and analogies can be drawn between these two designs. ^12,14^ In this study, the RPM was varied to enable it to cut a bone substitute to a precise particle size; lower RPM results in a larger particle size. Likewise, trephines at lower RPM produce larger particles.^25^ In two studies, the particle size was greater than1000□μm^7^ and 500 μm^25^ at 100 & 800 RPM respectively.

Though the particles were not spherical, there was no significant difference between length and breadth of the particles in both the groups. Particle shape, size and surface topography directly affect mechanical resistance to handling and compressive forces and play an important role in initiating cellular responses conducive to bone formation.^6,26^ A major concern is that Hydroxyapatite decomposes if exposed to high temperatures affecting its biocompatibility and resorption rate when used as a graft.^9,27^ Hence, the biocompatibility of particles obtained from both the groups was evaluated in an animal model. The mineralized tissue volume values were lower for SS & MS (68 & 66%) groups when compared to control (89%; autogenous core). Autogenous grafts integrate faster, and the volume of bone seen for SS and MS groups at 2 months was seen within one month for particulate autogenous graft in one study.^28^ However the values obtained in this study were slightly lower and higher to those observed by Takauti^29^ (78%) and Sadeghi^30^ (23%) in rabbits at 2 and 3 months for ceramics and nano-hydroxyapatite respectively. The possibility of thermal damage to the material appears remote in this study.

This study has some limitations worth noting; The milling chambers in commercially available laboratory milling systems are cooled with dry-ice or liquid nitrogen.^9,12,14^ We used neither; this may not have affected overall results as the device was used at a far lower RPM than above. Histopathology as applicable to rabbit bone defect model for basic biocompatibility testing was performed. Extensive histological evaluation of the bone healing was not the goal of this study.

To conclude, we report on the design and development of an automated milling system capable of generating particles of two different sizes. The device was able to generate small and medium-size graft particles which were distinct from each other in measures of particle count, diameter, distribution and size. The device can be adapted for a myriad of materials and protocols and aims to be a promising addition to the bone graft armamentarium.

## ACKNOWLEDGEMENTS

The authors wish to thank Dr Ramanand Oruganti for his valuable help with Stereomicroscopy. The authors declare that there is no conflict of interest.

## Notes

### Competing Interest Statement

The authors have declared no competing interest.

